# Independent Tryptophan pathway in *Trichoderma asperellum* and *T koningiopsis*: New insights with bioinformatic and molecular analysis

**DOI:** 10.1101/2020.07.31.230920

**Authors:** Uribe Bueno Mariana, JL Hernández-Mendoza, García Carlos Armando, Ancona Veronica, Violeta Larios-Serrato

## Abstract

The synthesis of Indole Acetic Acid from tryptophan has been described in plants, fungi and bacteria; it is thus known as tryptophan-dependent indole acetic acid. Four possible pathways of IAA formation have been described, including the indole acetonitrile acid (IAN), indole acetamide (IAM), indole-pyruvic (IAP) and tryptamine (TRM) pathways. Of these, the indole acetonitrile pathway is particularly important because when this compound is transformed into IAA, a nitrogenated molecule is released. The microorganisms that have this pathway are thus called nitrogen fixers. There is another little-studied pathway called TRP-Independent, so-called because the IAA that is formed in it can have an exogenous origin, chorismic acid (CHA), and it enters the pathway through anthranilic acid (ANA). The TRP-Independent pathway is made up of three stages. The first from CHA to ANA, the second from AA to IAA and the third from TRP to ANA through Kynurenine (KYN). This work describes the different stages of the pathway, as well as the enzymes and the genes that control the production of IAA, using a bioinformatic analysis of the genes involved, which were identified by PCR. An expression analysis showed that only *T asperellum* has the necessary genes to incorporate ANA into the TRP-I pathway and synthesize IAA through it. The analysis also detected the gene that regulates anthranilate phosphoribosyl transferase (AFT), an enzyme necessary for the synthesis of AIA from ANA; the presence of this gene was confirmed in the two species analyzed.

## BACKGROUND

Organisms of different phyla possess genetic tools for the synthesis of auxins. In plants, this mechanism allows them to produce growth-stimulating substances, but in other organisms such as bacteria and fungi, auxins do not play the same role. In plants IAA pathways that originate from tryptophan have been described, which is why they are called TRP-Dependent. These pathways involve tryptamine, indole acetamide, indole acetonitrile and indole pyruvic acid. Of these pathways, the indole acetonitrile pathway is relevant because when this compound is transformed into IAA, a nitrogen molecule is released into the soil in what constitutes a form of biological fixation of nitrogen.

*Trichoderma* spp. promotes plant growth through the production of auxin compounds (Dewick, 1998, Hoitink, *et al*., 2006; Vinale *et al*., 2008; Quittenden *et al*., 2009; Azarmi *et al*., 2011; Mano and Nemoto, 2012). Several studies have reported the synthesis of indole acetic acid (IAA) via tryptamine (TRM) and tryptophol (TIF) from tryptophan (TRP) (Gravel *et al*., 2007). The TRM pathway is part of the IAA synthesis pathways that have been described in plants, bacteria and other organisms (Aguilar-Piedras *et al*., 2003; Quittenden *et al*., 2009; Azarmi *et al*., 2011; Mano and Nemoto, 2012). There are reports of the synthesis of IAA and other auxinic compounds in *T atroviride, T viride* and *T harzianum*, but there is no information on the metabolic pathways or mechanisms used by a fungus alone or in association for the synthesis of the metabolite (Gravel *et al*., 2007; Harman *et al*., 2004).

A little known IAA synthesis pathway in plants, bacteria and fungi involves kynurenine (KYN) and is considered TRP-Independent (TRP-I). In this pathway, TRP is transformed into N-Formyl-kynurenine (NFK) by the action of the kyn A gene. Then it transforms into L-KYN by the action of kynurenine formidase (kyn B). Depending on the degradation pathway, several compounds can be formed from L-KYN, including kinuric acid by the action of kynurenine aminotransferase (2.6.1.7). Another pathway that starts with L-KYN, is the one that synthesizes 2-amino-3-carboxymuconate semialdehyde, from which picolinic acid or quinolinic acid can be produced. The TRP-KYN pathway has been widely studied in humans and other animals, since alterations of KYN metabolites are associated with neuronal disorders and even mite parasitosis (Krause *et al*., 2011; Kolodziej *et al*., 2011; Kolodziej, 2013; Ito *et al*., 2004).

In the case of *C psittaci*, Flavobacteriia and Xanthomonas, KYN is transformed into anthranilic acid (AA) by the action of kinureninase and is then transformed into 5-phosphoribosyl anthranilate, carboxyphenyl amino-1-Deoxyribulose-5-phosphate, and indole-3-glycerol phosphate, a direct predecessor of indole acetic acid. In this pathway, AA can also have an exogenous origin, derived from shikimic acid through chorismic acid. This pathway has also been described in Lima *et al*. (2009).

Four metabolic pathways for IAA have been reported in *Azospirillum brasilense* and other bacteria; these are the same pathways described in plants as TRP-D. The most important is the pathway involving indole acetonitrile, which, when transforming into indole-3-glycerol phosphate, releases a nitrogenated molecule. In *Azospirillum brasilense*, this mechanism is responsible for the biological fixation of nitrogen (Patten, 1996; Aguilar-Piedras *et al*., 2003).

An alternate degradation pathway of TRP leading to IAA has been described in *Streptomyces coelicolor*. This pathway, which involves the participation of Kynurenine (KYN), anthranilic acid and glycerol-3-phosphate indole, is coordinated by the genes SCOO3646, SCO3644 and SCO3645, which are homologous to the sequences reported for the genes kyn A, kyn B and kyn U (Lima *et al*., 2009; Zummo *et al*., 2012).

In the TRP-I pathway, TRP can synthesize IAA, passing through Kynurenine (KYN) and AA. This pathway has also been described in Flavobacteriia and Xanthomonas (Lima *et al*., 2009). Interestingly, the degradation of TRP through the KYN pathway into anthranilate has been described in humans; from anthranilate, it can pass to kynurenic acid, quinolinic acid or picolinic acid. Disturbances in this pathway can, in the case of humans, be associated with nervous disorders, injuries or psoriasis (Kolodziej *et al*., 2011; Kolodziej, 2013; Ito *et al*., 2004). It has also been found that TRP can be synthesized from 3-indole pyruvic acid in humans (Richards *et al*., 1972).

Apparently, IAA is stored in the body only in small quantities because when larger amounts are present, the body associates it with indole butyric acid, sugars or amino acids (Aguilar-Piedras *et al*., 2008). In fact, IAA is a direct precursor of AIB, which is formed when IAA passes through the microsomal membrane of corn cells, with acetyl CoA and ATP as cofactors.

In view of the scarce information available about the TRP-I pathway in *Trichoderma* spp, this work performed a bioinformatic and molecular analysis of the ability of *Trichoderma asperellum* and *T koningiopsis* to exogenously synthesize IAA through the TRP-independent pathway in liquid culture media with and without Tryptophan.

## MATERIAL AND METHODS

This study used cultures of *Trichoderma asperellum* (NRRL50191) and *T koningiopsis* (NRRL50190) isolated from the rhizosphere of *Helianthus annus* and from soil used for the cultivation of sorghum in the region where the state of Tamaulipas, Mexico, borders with Texas. For the HPLC, molecular and expression analysis, *T asperellum* and *T koningiopsis* were cultured in YPD medium with and without 100 ppm of TRP or IAA. Three repetitions per treatment were used, plus the control (PDB culture medium). The flasks were incubated at 25 °C for 96 hr and constant stirring at 200 rpm. Three mL per treatment and repetition were collected every 24 hr, centrifuged at 4500 rpm for 10 min and frozen at -20 °C until analysis.

### Bioinformatic analysis

A bibliographic review was made to establish the genes and enzymes of bacteria, fungi, plants and humans that have been reported to participate in the Trp-I pathway, since no study has made a complete description of this pathway. Nine genes were identified. The sequences and abbreviations with which they are reported change with the organism in which they were studied. This information was used to make a table (Table 1) detailing the name of the enzymes and genes corresponding to each organism, and their associated functions. The functions of enzymes and genes were established by searching the protein family database using multiple sequence alignment and Hidden Markov Models (PFAM). The available sequences were downloaded in the Stockholm file format. Subsequently, the hmmsearch function was used to compare the profiles found in the database with each of the proteins obtained from *T asperellum* and *T koningiopsis*, identifying the sequences homologous to each gene, and the statistical probability of sequence similarity. When the probability value exceeded 1 × 10^−3^, it was considered a significant result.

**Table 1.**
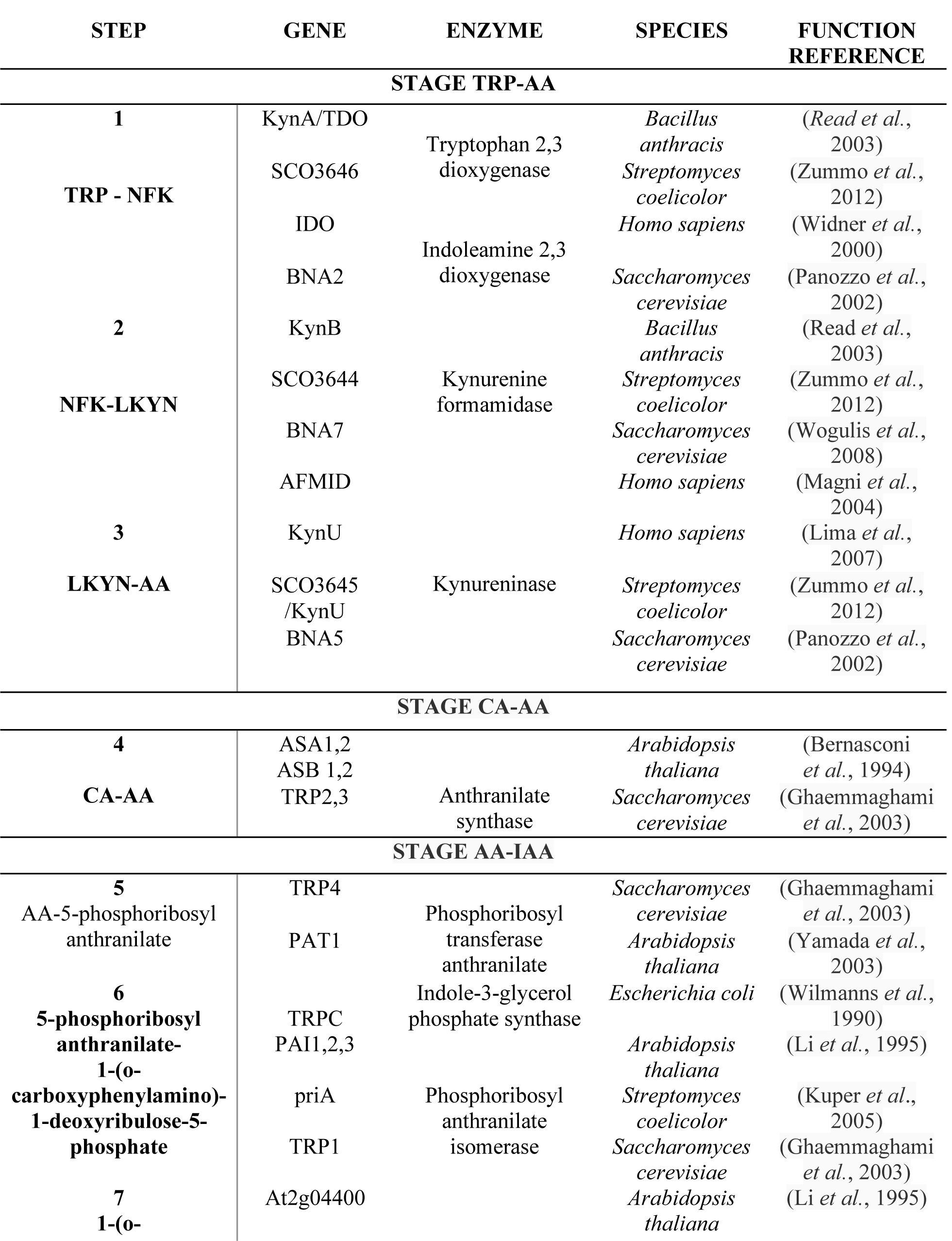
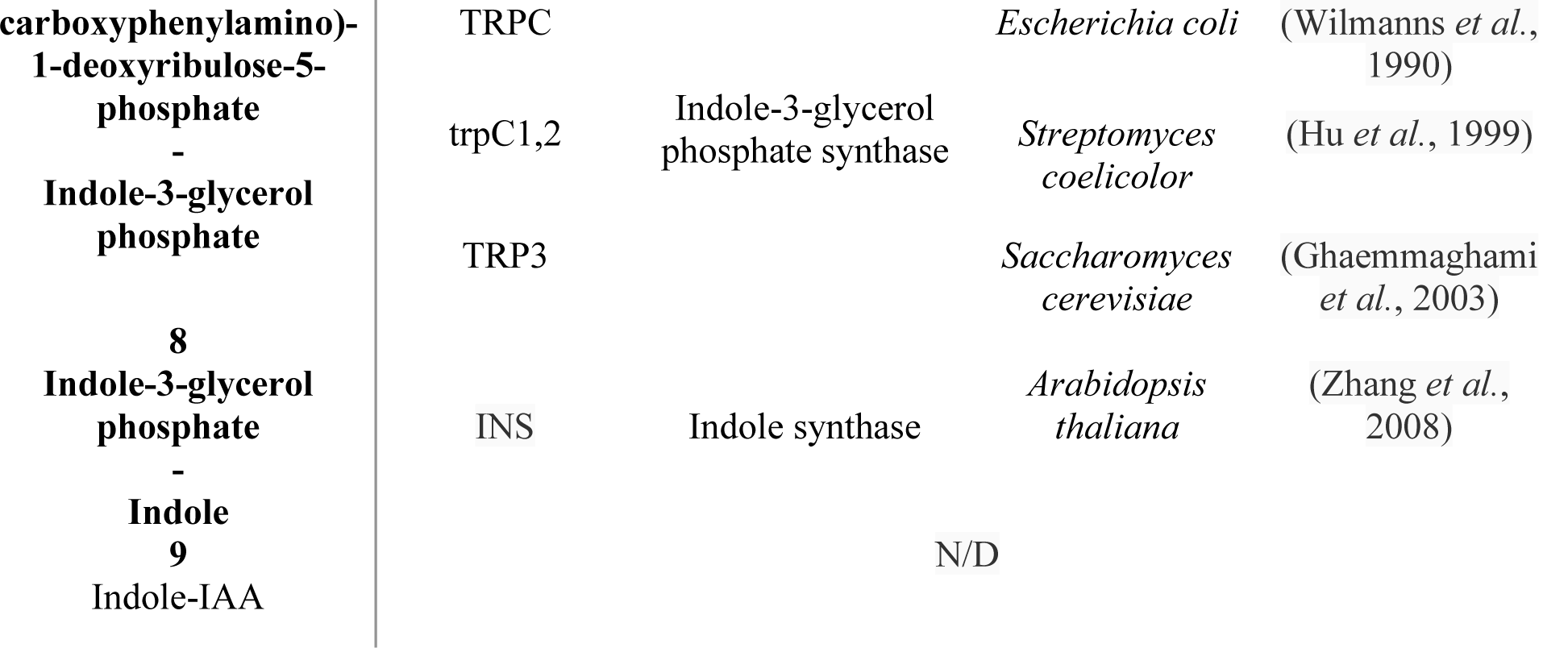
Enzymes/genes involved in the different steps of the TRP-Independent pathway in different organisms.

### Search for enzymes/genes participating in Trp-Independent pathways

The search for genes/enzymes of *Trichoderma* spp was carried out using the Mycocosm database of the Joint Genome Institute (JGI), belonging to the US Department of Energy. Protein sequences from the following 7 species of *Trichoderma* were downloaded: *T asperellum, T koningiopsis, T longibacterium, T citrino, T atroviride, T reesei, T harzianum* and *T virens*. A bioinformatic analysis was performed on these sequences.

### HPLC analysis

The samples were filtered through 0.45 μm nylon membranes (Millipore™; Cork, Ireland) and injected in HPLC (Hewlett Packard-Agilent™ model 1100), using a C18 ultrasphere column of 150*4.6 MM (Beckman Ultrasphere™, Fullerton, USA). The phase used was 80/20 (water-acetonitrile), pH 3, with a flow of 1mL/min, wavelength of 220 nM, using a UV detector (G1314A, Hewlett-Packard Agilent™). HPLC-grade products of tryptophan (TRP, Sigma-Aldrich), indole acetic acid (AIA, Fluka™), kynurenine (KYN, Aldrich™) and anthranilic acid (AA, Aldrich™) were used. All samples were run in triplicate (Hernández-Mendoza *et al*., 2010).

## RESULTS

### Bioinformatic analysis

This analysis showed that the AS gene, which regulates the synthesis of ANA from CHA, is present in the species analyzed using multiple sequence alignment and Hidden Markov Models (PFAM). No species of *T koningiopsis* have been sequenced; thus, the presence of this gene could not be determined in the strain 50190 studied here. The same is true for anthranilate phosphoribosyl transferase, which regulates the passage of ANA to 5 phosphoribosyl anthranilate; it is present in *T asperellum* but there are no data to suppose its presence in *T. koningiopsis*. This is also the case of ANA, which can be synthesized from L-Kyn with the participation of kynureninase. The Table 1 shows a summary of the bioinformatic analysis of this pathway, showing each enzyme of species with released genome, as well as the statistical value used to suppose its presence. The results of this analysis were obtained with the HMMER program.

For indoleamine 2, 3 dioxygenase, the value with the lowest chance of being chosen at random was 8.5e-166, corresponding to *Trichoderma atroviride*; the results of the 6 remaining species showed values close to that. It was found that T *asperellum, T virens, T harzianum*, and *T atroviride* possess the gene that codes for kynurenine formamidase (BNA7, SCO3644, KynB, AFMID) and that it is absent in the species *T citrino, T reesei* and *T longibrachiatum*. In the analysis for kynureninase, the best value found was 5.5e-136, corresponding to *T citrino*. Except for *T longibrachiatum*, the rest of the species had similar e-values. In the case of kynurenine formamidase, *T virens* had the best result, with an e-value of 2.7e-16. *T virens* also had the most significant e-value for anthranilate phosphoribosyl transferase (Table 4 in supplementary materials).

In the analysis for kynureninase, the best value found was 5.5e-136 in *T citrino*. Except for *T longibrachiatum*, the other species had similar e-values. In the case of kynurenine formamidase, *T virens* had the best result, with an e-value of 2.7e-16. *T virens* also had a significant e-value for anthranilate phosphoribosyl transferase.

Regarding anthranilate synthase, the best value was obtained in *T reesei*, with an e-value of 2.2e-26. *T virens* also had the highest e-value for phosphoribosyl anthranilate isomerase, with 7.7e-49. Indole-3-glycerol phosphate synthase was also analyzed; the best e-value (4.4 e-94) was found in *T virens*.

Significant and expected results were obtained when predicting protein structures using the HHpred tool, based on Hidden Markov Models. With respect to the sequence with code 100781 belonging to *T asperellum*, which was expected to code for an anthranilate synthase, an enzyme responsible for the transformation of chorismic acid into anthranilic acid, a 100% probability was recorded, as well as an e-value of 2e-111 for local prediction and of 2e-160 for global prediction. An image was taken of the superposition (Figure 1A in supplementary materials) of the predicted structure with the most representative structure, with an RMSD deviation (an indicator of divergence in aligned structures) of 0.653 A. Similar structures typically have RMSD values between 1 and 3 A; the higher the RMSD, the lower the similarity.

**Figure 1.**
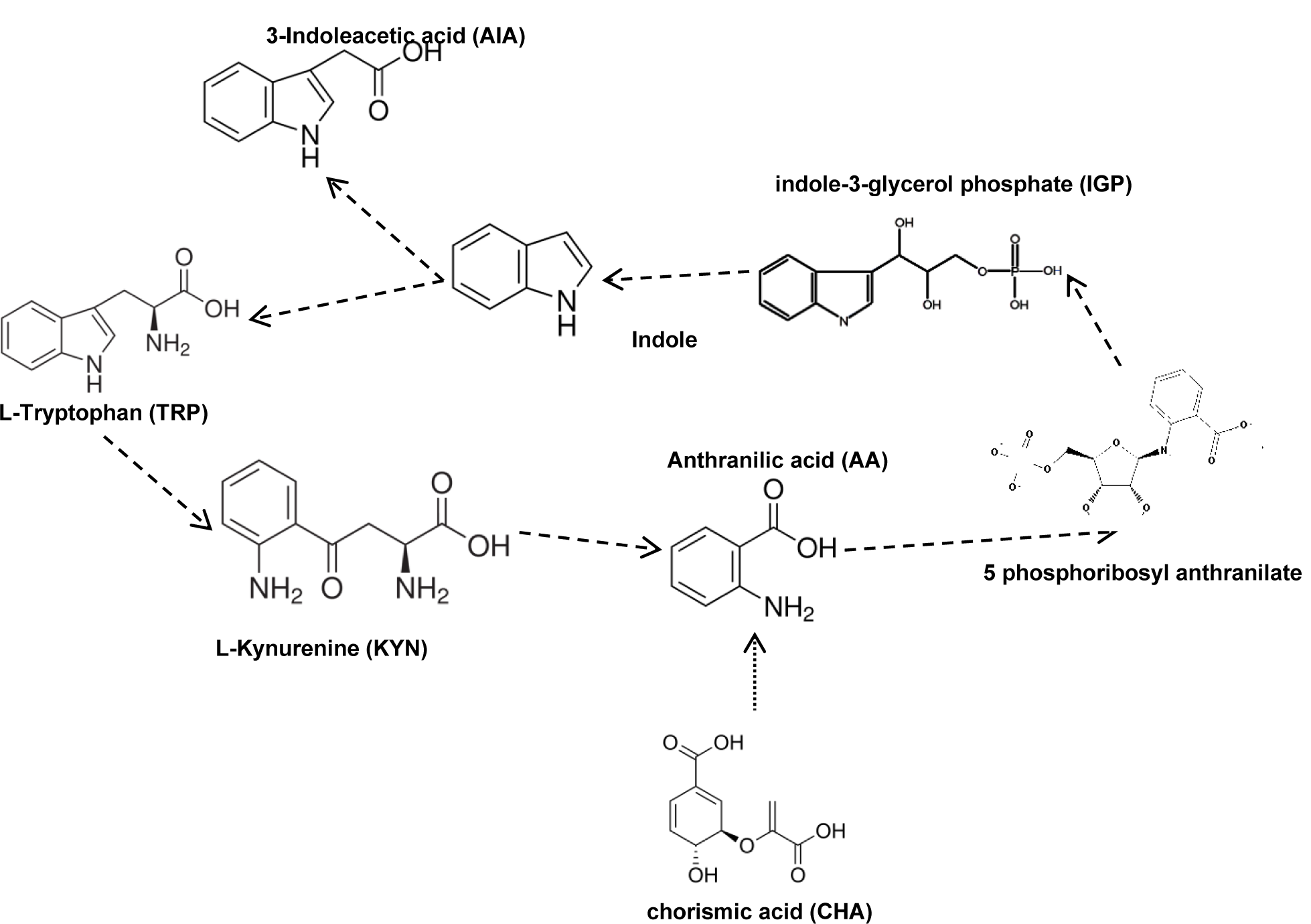
Proposed metabolic TRP-I pathway derived from bioinformatic analysis and HPLC analysis of culture medium with *T asperellum* and *T koningiopsis*. For *T koningiopsis* the anthranilate synthase (CHA→ANA) no is present, and this pathway of TRP-I don’t is present.

As for AS, an analysis of the structure of the anthranilate phosphoribosyl transferase protein was performed. This protein is involved in the transformation of anthranilic acid into 5-phosphoribosyl-anthranilate. The results showed that it had a structural probability of 100% of being the expected protein, with an e-value of 2.2e-89 for local prediction and of 2e-121 for global prediction. When the structures were superposed (Figure 1B supplementary materials) the RMSD deviation was 0.496 A, indicating a high degree of structural conservation.

## HPLC ANALYSIS

The analysis used to detect metabolites involved in the TRP-I pathway in YPD medium detected IAA in *T asperellum* after 96 h, whereas in *T koningiopsis* it detected it after 24 hr. Similarly, KYN and ANA were found in *T koningiopsis* in higher quantities than in *T asperellum*, which could indicate that this pathway is more active in *T koningiopsis*.

The detection of Tryptamine (TRM) and Indole Acetamide (IAM), which corresponds to the TRP-D pathway, was incorporated to verify the existence of other pathways for the synthesis of IAA in both species. These metabolites had already been detected in these species in previous works (unpublished). In the present study, the pathway involving TRM was more active in *T asperellum* than the IAM pathway, as evidenced by the higher concentrations of TRM (Table 2). In contrast, both metabolites were produced in similar concentrations in *T koningiopsis* (Table 3).

**Table 2a.** Metabolites produced by *T asperellum* in YPD culture medium with tryptophan as growth cofactor.

| Metabolite | 0 h | 24 h | 48 h | 72 h | 96 h |
| --- | --- | --- | --- | --- | --- |
| IAA | 0 | 0.53±0.28 | 0.21 ±0.25 | 0.26 ±0.22 | 0.27 ±0.47 |
| AA | 0 | 3.83 ±0.42 | 3.74 ±0.14 | 1.86 ±1.86 | 1.11 ±0.32 |
| TRP | 103 | 131.37 ±3.07 | 133.77 ±0.39 | 123.74 ±18.19 | 128.35 ±37.69 |
| TRM | 0 | 0.90 ±0.09 | 0.95 ±0.09 | 0.53 ±0.48 | 0.50 ±0.52 |
| KYN | 0 | 90.53 ±3.87 | 90.22 ±1.89 | 81.38 ±10.63 | 89.94 ±38.59 |
| IACM | 0 | 0.39 ±0.05 | 0.35 ±0.005 | 0.37 ±0.01 | 0 |

**Table 2b.**
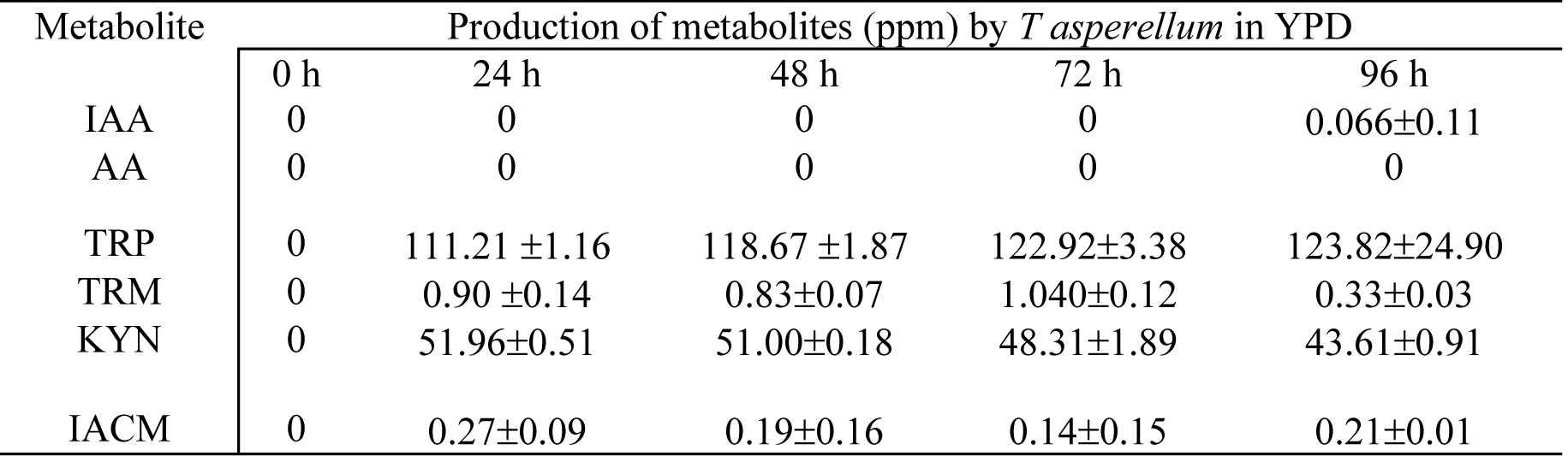
Metabolites produced by *T asperellum* in YPD medium with and without 100 ppm of TRP.

**Table 3.** Metabolites produced by *T koningiopsis* in YPD medium with and without addition of 100 ppm of TRP.

| Metabolite | Production of metabolites (ppm) by *T koningiopsis* in YPD + TRP |  |  |  |  |
| --- | --- | --- | --- | --- | --- |
|  | 0 h | 24 h | 48 h | 72 h | 96 h |
| **IAA** | 0 | 0.25±0.23 | 0.37±0.09 | 0.72±0.22 | 0.22±0.38 |
| **AA** | 0 | 1.09±1.88 | 1.75±1.52 | 2.68±0.33 | 2.88±2.54 |
| **TRP** | 0 | 173.16±61.14 | 270.51±77.21 | 164.64±82.34 | 173.34±63.04 |
| **TRM** | 0 | 0.85±0.41 | 0.63±0.23 | 0.53±0.03 | 0.46±0.39 |
| **KYN** | 0 | 85.09±7.17 | 98.01±13.18 | 74.13±29.92 | 74.57±9.66 |
| **IACM** | 0 | 0.88±0.17 | 0.63±0.16 | 0.18±0.03 | 0.47±0.30 |

| Metabolite | Production of metabolites (ppm) by *T koningiopsis* in YPD |  |  |  |  |
| --- | --- | --- | --- | --- | --- |
|  | 0 h | 24 h | 48 h | 72 h | 96 h |
| **IAA** | 0 | 0 | 0 | 0 | 0.49±0.30 |
| **AA** | 0 | 0 | 0 | 0 | 1.59±0.37 |
| **TRP** | 103.2 | 133.52±38.34 | 124.33±42.17 | 85.15±4.53 | 84.43±9.38 |
| **TRM** | 0 | 5.79±5.65 | 1.52±0.42 | 0 | 0.82±0.30 |
| **KYN** | 0 | 22.64±10.42 | 28.51±1.86 | 34.00±5.25 | 35.21±2.03 |
| **IACM** | 0 | 0 | 0 | 0 | 0.15±0.05 |

**Table 3.**
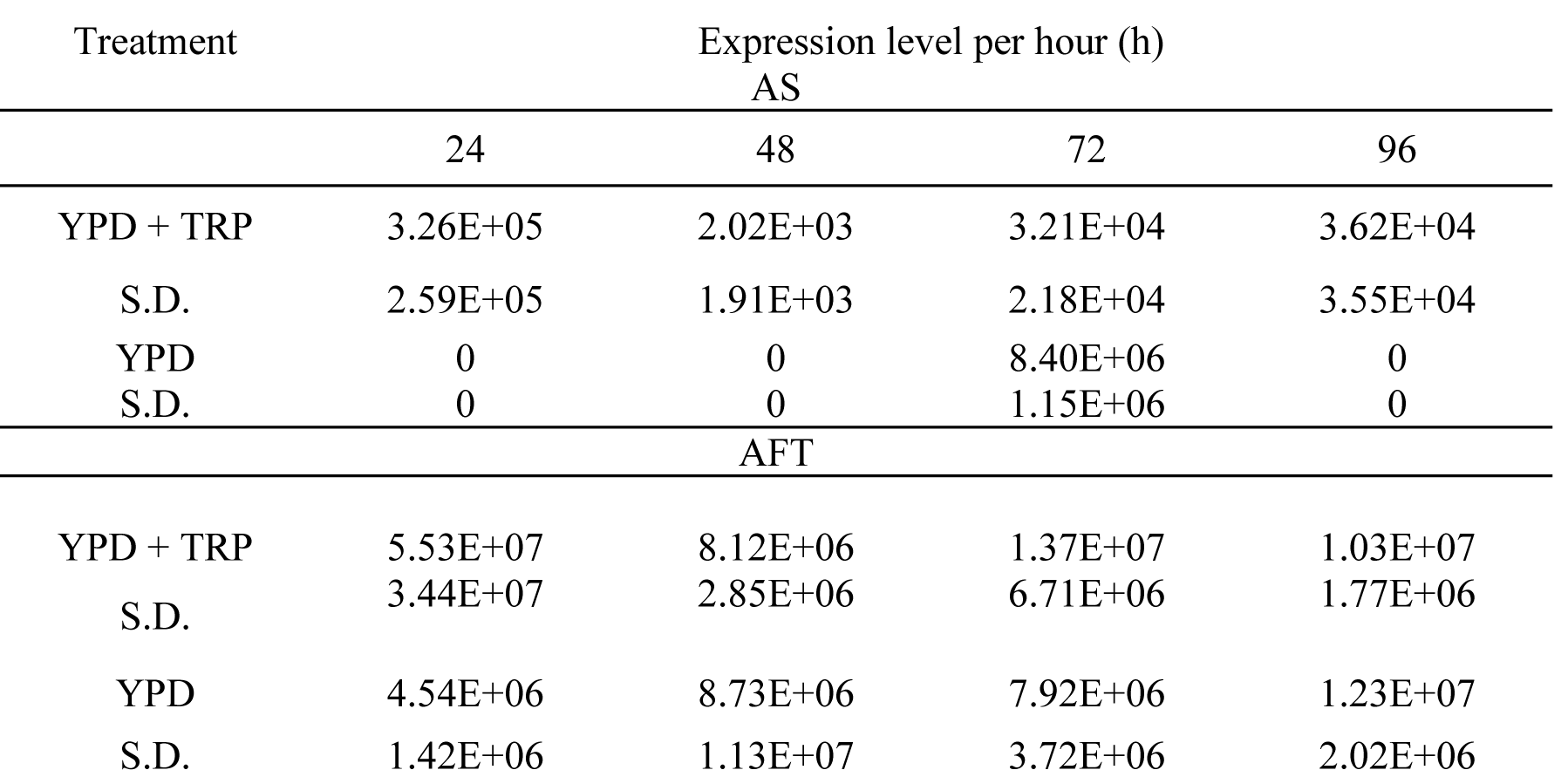
Average results of the intensity of expression in different treatments and times for the AS and AFT coding genes in *T asperellum*.

The addition of TRP to the YPD medium caused changes in metabolism; in *T koningiopsis*, IAA, ANA and IAM were detected only 96 hr after the start of the assay, while in *T asperellum* they were detected during the entire assay (Table 2). With respect to IAM and TRM, which participate in the TRP-D pathway, they were detected during the entire assay. The TRM pathway is apparently an active one, since TRM concentrations were higher than IAM concentrations. The auxin kynurenine was detected on another work using GC (no published)

### Amplification of the genes coding for AS and FRT

The primers for anthranilate synthase (AS) and anthranilate phosphoribosyl transferase (AFT) were evaluated using the *in* silico simulation platform of Molecular Biology Experiments of the University of the Basque Country. A fragment of 516 bp was obtained for AS and one of 176 bp for AFT. The PCR reactions for *T asperellum* generated individual bands of expected size for AS and AFT. It is worth noting that the PCR performed for AS in *T koningiopsis* yielded no product and that only the product corresponding to AFT was recovered. All the bands obtained were sequenced by Eurofins Genomics. The sequences were visualized with the computer program 4peaks and compared with NCBI data, which confirmed that they corresponded to AS and AFT. This information allowed to determine that the strain 50190 of *T koningiopsis* does not have a regulator for AS and is thus not able to obtain AA from CHA, which means that this pathway is not present in this strain.

### Analysis of gene expression for AS and AFT in *T asperellum*

RNA was extracted (in triplicate) from both strains at the same time for each sample (24 to 96 Hr) in order to perform the HPLC analysis. Figure 4 (in supplementary materials) shows the RNA extraction runs for *T asperellum* at 24 and 48 h (A), and at 72 and 96 h (B), in the treatments without TRP. (C) and (D) correspond to the same treatments, but with TRP added to the medium. The samples taken at 0 h were not subjected to RNA extraction since the amount of biomass was not enough to perform the analysis at that time, and the fungi had not started their growth and use of TRP.

### RT-PCR analysis of the gene encoding AS and AFT and quantification of transcript levels through the intensity of gene expression

We measured the transcript expression of the genes encoding anthranilate synthase (AS) and anthranilate phosphoribosyl transferase (AFT) in *T asperellum*. As control, 18S ribosomal (ITS1 and ITS4) and the oligos designed for AS and AFT, used to identify the respective genes. The amplified products of AS, AFT, and of the control gene 18S for T. asperellum, had fragments of 516 bp for AS, 176 bp for AFT and 632 bp for the control gene 18S. In the case of AS in treatment with TRP, fragments of the expected size were amplified in all samples, while in the medium without TRP (Figure 6 in supplementary materials), the only evidence of expression was recorded at 72 h. With respect to the control gene 18S, fragments of the expected size were amplified at all hours and all repetitions, both in the treatment with TRP) and in the treatment without TRP.

The results show that the addition of TRP strongly influences the expression levels of the regulatory gene of AS, since it appeared to be active throughout the entire sampling time in the medium with TRP. In the medium without TRP, by contrast, the gene was only detected at 72 hr. In the case of the AFT gene, the highest level of expression was recorded in the treatment with TRP at 24 h; in the treatment without TRP, the highest expression value was recorded at 96 h. The student’s t test showed that there were no differences (P=0.075 and 0.88) in the average expression values of the genes coding for AS and AFT between the treatments with 100 ppm of TRP and with no TRP.

## DISCUSSION

### Bioinformatic analysis

It is common to perform sequence homology searches using tools such as BLAST; however, beyond sequence coincidences, it is not possible to infer other aspects such as the function that corresponds to each sequence. Therefore, bioinformatic resources that are based not only on coincidences but on statistical aspects of the sequences have gained popularity, providing a much more useful alternative for inferring protein functions (Söding *et al*., 2005).

Only 1 of the 2 *Trichoderma* species studied in this work has been sequenced in its entirety. The amino acid sequences of *T asperellum* and those of other 6 species were used to minimize the variation derived from not knowing the sequences of one of the species of the study (*T koningiopsis*). These sequences were analyzed using Hidden Markov Models (HMMER). The 7 genes that participate in TRP-I pathways were analyzed; 6 of them, indoleamine-2,3-dioxygenase, kynureninase, anthranilate phosphoribosyl transferase, anthranilate synthase, phosphoribosyl anthranilate isomerase and indole-3-glycerol phosphate isomerase were present in all species analyzed, while kynurenine formamidase was absent only in *T citrino, T reesei* and *T longibrachiatum*.

The results obtained show that one of the key enzymes in the TRP-I pathway for the synthesis of IAA, anthranilate synthase, has very similar probabilistic values (e-value) between species, which could indicate a certain degree of conservation in this step. It can be inferred, at least bioinformatically, that the species studied in this work are able to produce IAA from TRP through a TRP-I pathway, since they show, with minimum error values, homologous sequences in their genomes that, although they do not are in the same probabilistic value ranges (e-value), always exceed the minimum set by the program (1×10^−5^) for reliable results. Analysis as this are not reported for *T asperellum* or *T koningiopsis*.

It is important to mention that the first step of this pathway involves both anthranilate synthase (AS) and anthranilate phosphoribosyl transferase (AFT), which are present, at least at bioinformatic level, in all *Trichoderma* species, which, in addition to those mentioned above, include *T virens, T atroviride* and *T harzianum, A thaliana, S cerevisiae, E coli* (Wilmanns *et al*., 1990; Bernasconi *et al*., 1994; Ghaemmaghami *et al*., 2003; Yamada *et al*., 2003) This type of analysis has not been tried before for *T asperellum* or *T koningiopsis*.

As it is known that between proteins with the same functions there are less differences in the structures than in the sequences, it was decided to analyze the structure of the AS and AFT proteins by means of a structural prediction tool based on the profiles from the Hidden Markov Models. The results showed that the two proteins had values of 2e-160 and 2e-121 respectively, corresponding to the global prediction model, which indicated that the length of the structure had been covered. Although up to now no studies have predicted the structure of AS and AFT in *Trichoderma* species, the importance of establishing an association between a molecular function and a protein structure has been addressed by Breda *et al*. (2008), who mention this technique as the best alternative to determine protein homology, as these proteins descend from a common ancestor and generally have the same structure and function.

The bioinformatic analysis is an excellent way to approach the search for genes and their presence in the genome of an organism, as in this case, the presence of the AS gen was rules out and thus the presence of this metabolic pathway in *T koningiopsis*. With this information, the proposal of the IAA synthesis pathway in *T asperellum* and *T koningiopsis* is made (Figure 2).

## CONCLUSIONS

The bioinformatic analysis performed with the published genomes shows that *T asperellum* contains the gene that coordinates the synthesis of AS, which controls the synthesis of anthranilic acid (ANA) from chorismic acid (CHA). It also has the gene that controls the transformation of ANA into phosphoribosyl anthranilate synthase, which means that it can also synthesize IAA from TRP through KYN. The HPLC and gene expression analyses confirmed this possibility.

In the case of *T koningiopsis*, there are not published genomes, so this analysis was not possible. The PCR performed on this fungus showed that the AS gene is not present. It follows that *T koningiopsis* (NRRL50191) does not have the genetic tools to synthesize IAA from CHA; however, it does have the kynureninase gene (reported in other works as SCO3645/KynU/BNA5), which controls the transformation of KYN into ANA, and the AFT gene, which ANA to transform into AFR and from there to IAA. This indicates that *T koningiopsis* can synthesize IAA from TRP via KYN.

## ACKNOWLEDGEMENTS

This work was financed by the Instituto Politécnico Nacional. Uribe-Bueno had a CONACYT scholarship for Master of Science studies. Hernández-Mendoza has COFAA, EDI-IPN and Sistema Nacional de Investigadores (SNI).

